# Metabolomic approaches highlight two mechanisms of accelerated grain filling in Mediterranean oat (*Avena sativa* L.) cultivars during drought

**DOI:** 10.1101/2023.06.28.546978

**Authors:** Aiswarya Girija, Francisco J Canales, Bahareh Sadat Haddadi, Rachel Dye, Fiona Corke, Jiwan Han, Jason Brook, Kevin Williams, Manfred Beckmann, Elena Prats, John H Doonan, Luis A J Mur

## Abstract

Grain filling in cereals is complex process that determines the final grain yield and quality. Abiotic stresses can have major impact on grain filling. Oats (*Avena sativa* L.) is sensitive to drought which adversely affect yield and productivity. In this study, we characterised the grain filling responses of two Mediterranean oat cultivars Flega and Patones under severe drought. Grains from the top (older) and bottom (younger) spikelets of primary panicle were larger in size in response to drought, particularly in Patones, suggesting accelerated grain development. The metabolomes of source (sheath, flag leaf) and sink (developing grains) tissues were profiled to describe source-sink partitioning. In Patones, the developing grains showed increased sugars and amino acids which indicate accelerated grain filling. These were associated with elevated α-linolenic acid levels in source tissues but decreased in developing grains under drought. There was also a significant decrease in C18 fatty acids (FA) and jasmonates (JA) derivatives in the developing grains which suggested a role for JA signalling in Patones with drought. Flega showed a different response, with accelerated flowering and enhanced energy metabolism in both source and sink organs. The accumulation of ophthalmic acid in grains of Flega and lower levels of reduced glutathione in source tissues suggested greater oxidative stress than Patones under drought may be driving the grain filling phenotype. This study suggests that oats cultivars can use α-linolenic acid-linked signalling or oxidative events influences accelerated grain filling with drought. These could be important traits in developing oat cultivars that maintain yield in drought-prone environments.

**Highlight:** The impact on drought in one tolerant and one susceptible oat cultivar was assessed at the grain filling stage. The drought tolerant cultivar, Patones, showed accelerated grain development which could be a strategy to escape drought. Metabolite mapping of flag leaves, sheath and grains of Flega suggested that alpha linolenic acid could be regulating the altered sink-source relationships. The drought susceptible cultivar, Metabolomics shifts in Flega suggested that oxidative stress accelerated flowering.

## Introduction

Grain filling is the most sensitive and important developmental stage that determines the yield potential of crops. Grain filling process is controlled by complex physiological, metabolite and cellular events and is directly and/or indirectly influenced by abiotic and biotic factors (Ma et al., 2023). Drought stress during grain filling leads to diminished photosynthetic activity which affects the mobilisation of nutrients from source to developing grains (Yan et al., 2022). This will drastically affect grain development and ultimately yield. In cereals, the mobilisation of nutrients from source is influenced by many factors, including genotype, environment, irrigation, fertilisation, plant density and photosynthetic ability (Tovignan et al., 2020). For example, the total weight in wheat and barley is determined by the grain filling rate, whereas in rice seed filling duration is more important and in maize the duration and rate are both important (Haverroth et al., 2021; Kennedy et al., 2018). However, despite the impact on yield, the key components that regulate grain filling need further characterisation.

In cereals, grain filling is determined by grain number and the size and final grain size is correlated to grain filling capacity (McCabe & Burke, 2021). This is further related to the remobilization of stored assimilates and photosynthetic carbon assimilation from vegetative tissues (source) to grains (sink) (Takahashi et al., 1993). Drought stress at post-anthesis stage alters photosynthetic efficiency and the re-mobilization of assimilates to developing grains from source (Shirdelmoghanloo et al., 2022). Grain number could also be reduced in wheat exposed to drought during reproductive development (Onyemaobi et al., 2016). In rice, drought enhanced starch accumulation in developing grains with increased ADP-glucose pyrophosphorylase (AGPase) and starch synthase (SS) activity. These studies suggest the importance of nutrient partitioning from vegetative tissues to grains that determine the final grain yield. Therefore, understanding the response of source and sink during drought is crucial for determining the grain yield and quality in cereals growing under semi-arid and drought-prone climates.

Oat (*Avena sativa* L.) is ranked sixth in global cereal production in terms of tonnage and is a dual-purpose crop for food and animal feed (Xie et al., 2021). Compared to other cereals, oat grains are a rich source of dietary fibre (β-glucan), protein (avenins), lipids, antioxidants (avenanthramides) and phenolics (tocols and saponins) (Allwood et al., 2021). Due to its nutritional and health benefits oats have gained attention in the food, pharmaceutical and cosmetic industry (Martínez-Villaluenga & Peñas, 2017). Compared to other cereals, oats are well-adapted to marginal environments, and tolerate marginal soils with low nutrient levels (Canales et al., 2021b; Kutasy et al., 2021). However, oat has high transpiration rate with relatively higher-water requirements than other cereals (Xiao & Yang, 1992) and are susceptible to grain abortion under drought (Kutasy et al., 2021). Therefore, drought is a major limiting factor in oats (Rispail et al., 2018; Wang et al., 2020).

The impact of drought stress during grain filling and the changes associated with source-sink flux have not been defined in oats. Here we apply untargeted metabolomics to investigate the grain filling performance of two contrasting Mediterranean oat cultivars (cv) ‘Flega’ and ‘Patones’ under drought. Flega is more susceptible to drought than Patones, which is relatively tolerant (Sánchez- Martín et al., 2018; Sánchez-Martín et al., 2014). This study evaluates the impact of drought on grain filling in oats using metabolomic approaches to characterise shifts in source and sink relationships. We found that with Patones in particular, oat plants showed accelerated grain filling, that could be influenced by α-linolenic acid. In Flega, the grain filling phenotype could be related to oxidative stress.

## Materials and Methods

### Plant Material growth condition and sampling

All experiments used two oat cvs Flega and Patones. Patones exhibits good adaptation to Mediterranean agroclimatic conditions and was developed by the ‘Instituto Madrileño de Investigación y Desarrollo Rural, Agrario y Alimentario’ (IMIDRA, Madrid, Spain). Patones was provided by the Plant Genetic Resources Centre (INIA, Madrid, Spain) whereas Flega was developed by the Cereal Institute (Thermi-Thessaloniki, Greece). The two varieties are genetically not closely related (Montilla-Bascón et al., 2013).

Plants were grown under controlled conditions on the LemnaTec platform at the National Plant Phenomics Centre (NPPC) (20°C, ambient relative humidity and under 14h /10h light/dark cycle with >450 μmol m^−2^ s^−1^). Plants (n=8 per experimental class) were sown in pots (15 × 15 × 20 cm) with one plant per pot in Levington F2 peat-based compost. Drought was imposed by automated watering to a reduced target weight at growth stage GS55 (panicle half emerged, (Zadoks et al., 1974) Plants were automatically watered to achieve a soil water capacity (SWC) of 25 % (severe drought) and 75 % (control) which was then maintained until harvest. The primary panicle of well-watered and drought plants was harvested on the 15^th^ day (GS75, grains at milky stage). The number of whorls and number of spikelets per whorl were recorded and the leaf-sheath (S), flag leaf (Fl), rachis (R) and spikelets of the primary tiller were sampled. Samples were flash frozen and used for metabolite extraction **(Fig. S1)**.

After sampling, the plants were maintained at the same relative soil water content and, upon maturity, the shoot biomass, total plant weight, number of tillers, number of stems, stem height, number of panicles, total panicle weight, number of whorls, number of spikelets, number of sterile spikelets, number of stems, panicles height and number of tillers were recorded.

### Stomatal conductance and Fv/Fm

F_v_/F_m_ was measured throughout the drought experiment using the Handy PEA+ system (Hansatech Instruments Ltd, UK). Stomatal conductance was measured in with an AP4 cycling porometer (Delta-T Devices Ltd, Cambridge, UK). The porometer was used on the mid of the adaxial surface of leaf laminae. Both stomatal conductance and F_v_/F_m_ was measured from the flag leaf (Fl) of the primary tiller (n=8).

### Metabolite profiling and statistical analysis

Freeze dried samples (20 ± 1 mg) were extracted in 500 uL of chloroform:methanol:water (1:2.5:1, v/v/v). An aliquot of 100 μL of the supernatant was transferred to an HPLC glass vial with a 0.2 mL micro insert (Kattupalli et al., 2021; Allwood et al., 2006). Untargeted metabolite profiling was undertaken by Flow injection electrospray mass spectrometry (FIE-MS) based on a Q Exactive plus mass analyser instrument with UHPLC system (Thermo Fisher Scientific©, Bremen, Germany). Mass-ions (*m/z*) were acquired in both positive and negative ionisation modes and data were normalised to the total ion count. Statistical analyses were performed on log_10_-transformed values. Multiple comparisons and post hoc analyses used Tukey’s Honestly Significant Difference (Tukey’s HSD) and Fisher LSD test (*P*< 0.05) to determine significant compounds. Principal component analysis (PCA), one-way analysis of variance (ANOVA), and hierarchical cluster analyses (HCA) used the online R-based platform, Metaboanalyst 5.0 (Pang et al., 2022). The significant hits were identified based on accurate masses (5 ppm resolution) and the ionisation patterns linked to that particular metabolite and associated isotopic forms. This involved comparison with the *Oryza sativa* japonica database in the Kyoto Encyclopaedia of Genes and Genomes (KEGG)(http://www.genome.jp/kegg/) and also the Human metabolite (HMDB), PubChem, and ChEBI databases.

## Results

### Effect of drought on physiological, developmental and yield parameters

Comparing the water usage data, both cultivars showed reduced total cumulative water use under drought period **(Fig. 1A).** Flega showed an overall significant reduction in the total water usage under both control and drought conditions as compared to Patones (**Fig. 1B)**. F_v_/F_m_ did not significantly change with drought treatment or between cultivars **(Fig. 2A)** but, flag leaf stomatal conductance showed a significant difference between the treatments but not between cultivars **(Fig. 2B)**.

**Figure 1.**
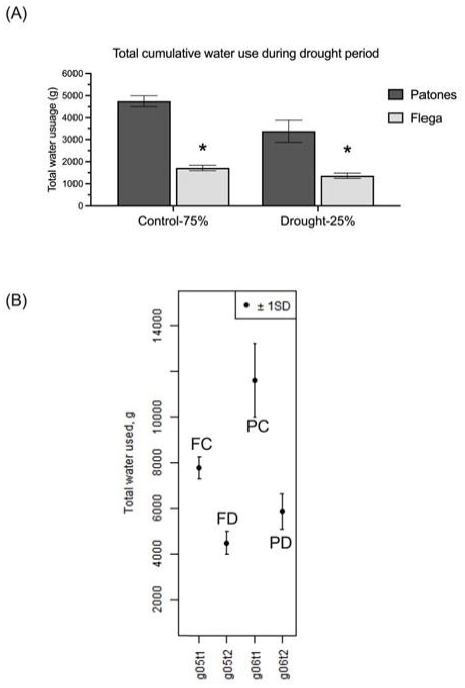
Cumulative water use data in Patones and Flega during the study. **(A)** Cumulative water used during drought period and **(B)** Total cumulative water during the whole experiment measured in Patones (P) and Flega (F) under control (C,75%) and drought (D, 25%) SWC. The data shown as average ± SD, n=8 and significance, *P* ≤ *0.05* based on Student’s t-test is shown by the black asterisk *. The data was acquired from the automated Lemnatec Platform.

**Figure 2.**
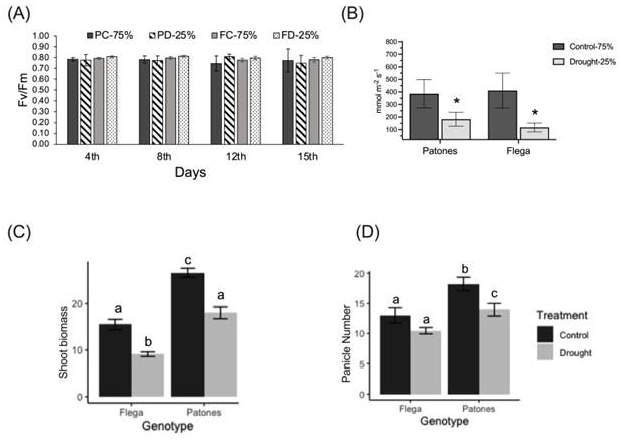
Morphological and physiological parameters measured during drought. **(A)** Fv/Fm was measured during the drought period. The data was collected periodically on day 4^th^,8^th^, 12^th^ and 15^th^. (**B)** Stomatal conductance from flag leaf of primary panicle measured in Patones (P) and Flega (F) under control (C, 75%) and drought (D, 25%) SWC. Morphological parameters collected on the day of sampling on from Patones and Flega **(C)** shoot biomass and **(D)** number of panicles. The data shown as average ± SD, n=8 and significance based on Student’s t-test P ≤ *0.05* is shown by different letters and black asterisk *.

Total dry shoot biomass, total plant weight, stem height, panicle weight, number of panicles, number of stem and number of tillers was measured at the end of the treatment. From these parameters, there were significant differences between control and drought treatments in both cultivars in shoot biomass, total plant weight, stem height, number of tillers, number of panicles, panicle height, panicle weight. **(Fig.2, Fig.S2).** Patones showed significant reduction in shoot biomass and number of panicles under drought, that were greater then with Flega **(Fig.2C, 2D)**. Flega showed an accelerated development and flowering a week earlier than Patones.

In oats, grain maturation begins at the panicle tip and proceeds towards the base. Within a spikelet the florets develop acropetally. The floret numbers can vary considerably (typically 1 to 4) in each spikelet, with grain filling rates correlated with spikelet and grain number (Haverroth et al., 2021). Therefore, a single snapshot of the grains on a given panicle can represent a developmental series that reflects grain maturation. Here we studied the phenotypes of developing grains in each whorl from the upper and bottom spikelet of the primary panicle and the grains were analysed for altered development in response to drought **(Fig.3)**. On examining the grain phenotypes, it was notable that drought led to larger grain sizes in both cultivars **(Fig. 3A).** This suggested accelerated grain development, which could be considered as an “escape” mechanism as it would allow earlier completion of the life cycle under drought. This feature was more prominent in each whorl of Patones and particularly in the youngest whorl (number 4) (**Fig. 3A, right panel)**. The number of spikelets in the basal whorls (3 and 4) was significantly higher (*P <* 0.05) in Flega compared to Patones **(Fig. 3B)**. To test the effect of treatment and cultivar on the metabolism of source and sink tissues, we undertook metabolomic analysis of different tissues including the flag leaf as a representative source and the developing grains as a sink.

**Figure 3.**
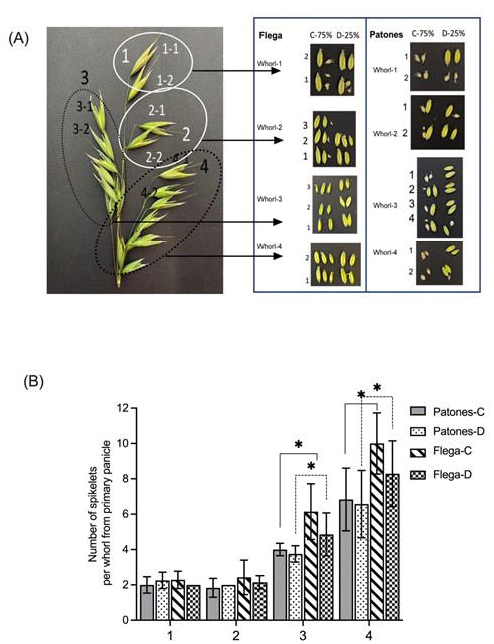
Grains collected from the primary panicle of Flega and Patones. **(A)** Primary panicle with four whorls, numbered from top to bottom. Top whorls 1 and 2; bottom whorls 3 and 4 with spikelets number showing 1 to 2. The corresponding grains separated from Flega (left) and Patone (right) from control and drought stressed plants are indicated using black arrows **(B)** Number of spikelets in each whorl in the primary panicle of Flega and Patones under control (C) and drought conditions (D). The data shown are average ± SD (n=8) with significance based on Student’s t-test, shown as *, *P* ≤ *0.05*.

### Drought induced metabolite changes in source (sheath, flag leaf and rachis) and sink (developing grains)

Source organs including the flag leaf (FL), sheath (S) rachis (R) and sink (developing grains) were harvested for nontargeted metabolite profiling. A total of 1947 and 2077 *m/z* were captured in negative and positive ionisation modes, respectively. PCA of all *m/z* features indicated altered metabolite profiles between source and sink tissues. Each cultivar showed drought specific responses with regard to each organ **(Fig. S3)** where the major sources of variation were linked to the organ across principal component (PC)1 but responses to drought were seen across PC2.

To understand the contribution of source and sink organs, further analyses were focused on tissue-specific metabolite profiles. With drought, the Flega sheath metabolomes showed a shift across PC2 whereas with Patones these showed a smaller response with a considerable overlap between the control and drought group clusters. **(Fig. 4A).** In the flag leaves, cultivar specific differences were prominent in the metabolome (across PC1), but drought responsive changes were seen in both cultivars across PC2 **(Fig. 4B)**. More prominent differences were observed in the rachis of both cultivars, under control and drought conditions **(Fig. 4C).** To obtain in-depth understanding of the key changes between the cultivars, pair-wise analysis was carried between control and drought samples for each cultivar.

**Figure 4.**
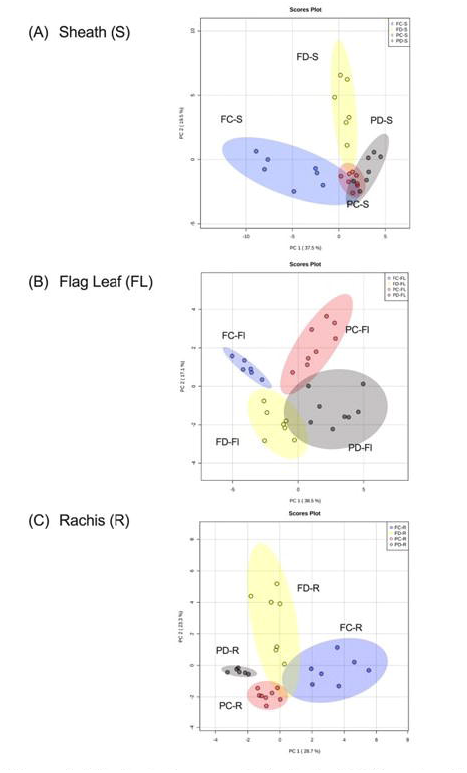
Principal Component Analysis (PCA) and pathway enrichment of metabolites. PCA mapping of annotated metabolites profile on the left panel and pathway enrichment on the right panel in different source tissues **(A)** Sheath (S), **(B)** Flag leaf (Fl) and **(C)** Rachis (R) of Flega and Patones.

Flega showed a greater metabolomic shift in response to drought in all the source organs **(Fig. 5)**. In this cultivar, the largest responses were observed in the sheath where pair wise comparisons of control (FC-S) and drought (FD-S) samples identified 201 significant *m/z* hits (**Fig. 5A)**. Assessment of these suggested that drought resulted in higher levels of TCA and glycolysis intermediates (citric acid, malic acid, pyridoxal 5’-phosphate, pyruvic acid), sugars (sucrose, raffinose, myo-inositol stachyose, rhamnose, deoxyribose,), organic acids (glyoxylic acid, hydroxypropionic acid, oxalic acid), gibberellin, behenic acid, quercitrin/kaempferol-3-O-glucoside and arachidic acid. However, myo-inositol-1-phosphate, gamma-aminobutyric acid (GABA), amino acids (glutamine, asparagine, aspartic acid, glutamic acid, lysine), glutathione (GSH), oxidized glutathione (GSSG) and N-acetyl-D-glucosamine phosphate were significantly reduced. In the flag leaf, the levels of 21 metabolites were found to be significantly different between control and drought treatment **(Fig. 5B)** with elevated levels of citric acid, hydroxypyruvic acid, aconitic acid, linolenic acid, glyceric acid, sulfate, raffinose, stachyose, pyruvic acid and palmitoleic acid in drought treatment. Ophthalmic acid **(Fig. 5B** indicated by blue*), a marker of oxidative stress in plants (Servillo et al., 2018) was also identified in Flega leaves under drought. This observation, along with the lower levels of reduced glutathione **(Fig. 5B** indicated by blue*) were consistent with different source organs and indicated an increased oxidative stress status in Flega. In the rachis, 25 features significantly differed between control and drought treatments **(Fig. 5C)**. The rachis exhibited drought increased accumulation of sucrose, citric acid, pyruvaledhyde, pyruvic acid, rhamnose, pyridoxal-5’ phosphate, keto-glutaramic acid, and tryptophan whereas it showed decreased levels of ribose, glutamate, asparate, alanine, glycine, the ethylene precursor 1-aminocyclopropanecarboxylic acid (ACC) and reduced glutathione **(Fig. 5C)**. Overall, Flega droughted plants, showed increased levels of citric acid, pyridoxal 5’ phosphate, stachyose, raffinose, and rhamnose but reduced levels of glutathione in the three source organs analysed. This suggests a shift in the redox-energy status leading to oxidative stress conditions in Flega under drought.

**Figure 5.**
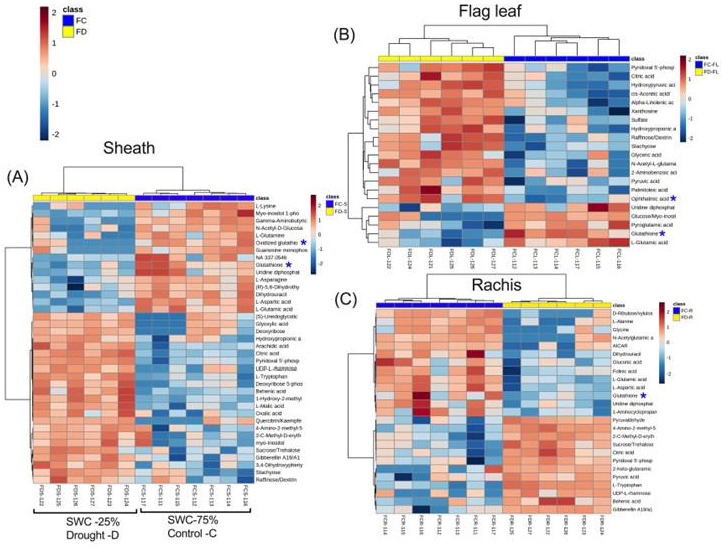
Metabolite distribution in different source organs of Flega. Heat maps showing the levels of significant metabolites in **(A)** Sheath-S **(B)** Flag leaves-Fl **(c)** Rachis- R of Flega (F) control (C, 75%) and drought (D, 25%) SWC. Blue asterisk indicates the levels of ophthalmic acid and glutathione in all the three source organs. Blue and red color in the heat map indicate low to high levels of features.

In Patones, the sheath exhibited changes in sugars, TCA/glycolysis intermediates and organic acids; namely, sucrose, citric acid, oxalic acid, glutaric acid pyridoxal 5’-phosphate and pyruvic acid were elevated in droughted group **(Fig. 6A)**. Flag leaf and rachis of Patones proved to have some distinctive responses to drought compared to Flega. There appeared to be similar changes in raffinose, stachyose and α-linolenic acid but others (ribose, gluconic acid, xanthosine, hydroxy pyruvic acid, hydroxy propionic acid, 2-keto-glutatamic acid, lauroyl-CoA, mercaptolactic acid) were only seen in Patones flag leaves **(Fig. 6B, C)**. In the rachis, there were common drought responsive changes in the levels of AICAR (5-aminoimidazole-4-carboxamide ribonucleotide, an intermediate in inosine monophosphate metabolism), L-tryptophan, L-aspartic acid, citric acid, pyridoxal 5’-phosphate, gibberellin, UDP-L-rhamnose, pyruvic acid and behenic acid in both Patones and Flega. **(Fig. 5C, 6C)**. Changes in α-linolenic acid were only seen in Patones and in all three source organs under drought **(Fig. 6D).** A drought induced decrease in reduced glutathione levels were seen in Patones sheath but not in the flag leaf and sheath. There were also no significant changes in ophthalmic acid in Patones. These observations could indicate that drought induced oxidative stress was not as severe in Patones compared to Flega.

**Figure 6.**
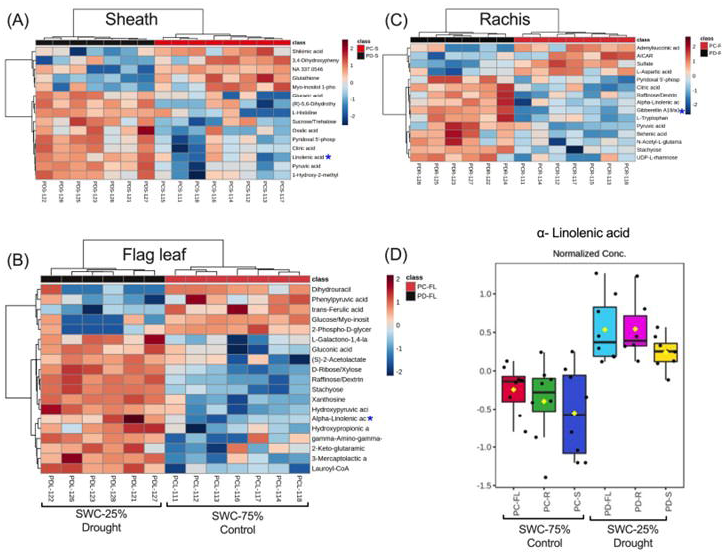
Metabolite distribution in source organs of Patones. Heat maps showing the levels of significant metabolites in **(A)** Sheath-S **(B)** Flag leaves-Fl **(c)** Rachis- R of Patones (P) control (PC) and drought (PD) plants. Blue and red color in the heat map indicate low to high levels of features. Blue asterisk indicates the levels of α-Linolenic acid in all the three source organs. **(D)** Box plot showing the levels of α-Linolenic acid in all source organs in Patones control (C, 75%) and drought (D, 25%) SWC plants.

### Drought induced metabolite changes in sink organs: Developing grains

To assess how developmental patterns were reflected in the sink metabolomes, developing grains devoid of lemma and palea were isolated from upper and lower whorls and analysed.

The developing grains from droughted Patones showed decreased accumulation of fatty acid derivatives including C18 metabolism and α-linolenic acid metabolism. In the older grains, drought resulted in a relative decrease in fatty acid related metabolites (linoleic acid, oleic acid, palmitic acid, palmitoleic acid, myristic acid, linolenic acid, arachidic acid) **(Fig. 7A)**. This pattern was also observed for 12-oxo-phytodienoic acid (12-OPDA), a precursor in jasmonate biosynthesis and a signalling molecule in inducing the expression of stress genes (Dave & Graham, 2012; Savchenko & Dehesh, 2014). Beyond changes in fatty acid related metabolites, endogenous levels of glutathione (GSH) and oxidized glutathione (GSSG) were found to be significantly higher in grains from both, upper and lower whorls, under drought stress. Younger grains, collected from the basal spikelets, showed more significant metabolite changes under drought than the older grains **(Fig.7B)**. These included the same relative decrease in fatty acids and 12-OPDA that was seen in the older grains **(Fig.7A and B, boxed)**. In addition, drought treatment in Patones increased the content of sugars, TCA and glycolysis intermediates, organic acids, and amino acids in younger grains **(Fig.7B)**. These changes could be associated with accelerated and efficient grain filling that resulted in a larger grain size as illustrated in **Fig.3**.

**Figure 7.**
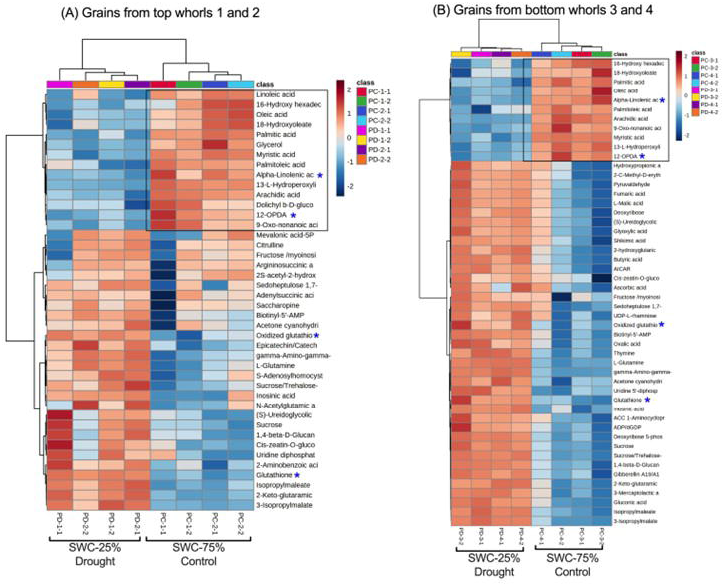
Metabolite mapping in Patones grains. Heat map showing metabolite levels in grains from Patones **(A)** grains from top whorl 1 and 2 **(B)** grains from bottom whorls 3 and 4 control (C-75%) and drought (D-25%) plants. The data shown are average of significant metabolites with n=8. Black box highlights the metabolite cluster associated with fatty acids and jasmonates metabolite pathways. Blue asterisk indicates 12-OPDA, α-Linolenic acid and glutathione levels.

Droughted Flega grains showed a distinct metabolite pattern to that observed for Patones, with no significant changes in fatty acid or jasmonate metabolism. The grains from both, upper **(Fig. 8A)** and lower spikelets (**Fig. 8B)** showed increased levels of citrulline, 3-isopropylmalate, O-phosphoethanolamine, saccharopine and O-acetyl-L-homoserine **(Fig. 8, Fig. S5)**. Apart from this cluster, apical grains showed increased levels of 2-keto-glutaramic acid and 1D-myo-inositol-1- phosphate. Interestingly, ophthalmic acid was targeted in the basal grains of both control and droughted Flega plants. Overall, the basal grains under drought showed an enhanced accumulation pattern for TCA and glycolysis intermediates, sugars, sugar phosphates and amino acids. Given that Flega showed earlier flowering that Patones, these metabolomic changes, possibly linked to oxidative stress, could contribute to a different ‘drought-escape’ mechanism.

**Figure 8.**
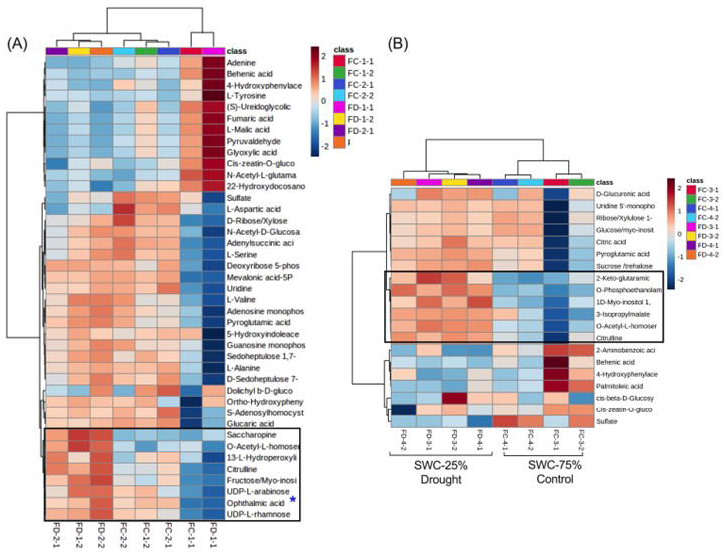
Distribution of metabolites in Flega grains. Heat map showing metabolite levels in grains from Flega **(A)** grains from top whorl 1 and 2 **(B)** grains from bottom whorls 3 and 4 control (C-75%) and drought (D-25%) plants. The data shown are average of significant metabolites with n=8. Black box highlights the metabolite cluster associated oxidative stress and energy status. Blue asterisk indicates the level of ophthalmic acid.

## Discussion

Drought is a crucial abiotic stress that affects crop production and represents a major global challenge to agriculture and food production (Ullah et al., 2022). Plants respond to drought by reprogramming physiological processes such as stomatal closure, transpiration rate, photosynthetic capacity, water usage. (Wu et al., 2017). Therefore, it is important to explore the molecular and regulatory networks that contribute to the drought tolerance in crops.

Grain filling is an important agronomic trait that determines the final grain quality and yield in cereals (Yan et al., 2022). The grain filling rate is also correlated to relative source-sink strength and nutrient partitioning (Shirdelmoghanloo et al., 2022). Compared to other cereals, oats have a higher transpiration rate and are sensitive to water deficit environments particularly at anthesis and grain filling, which typically result in grain shrinkage and yield reduction (Zhang et al., 2022). There is limited knowledge on the impact of drought on grain filling in oats, so here we characterised the drought induced responses on the metabolomes of sink and source tissues in two contrasting cultivars; Flega and Patones whose drought responses had been previously characterised at the seedling stage (Sánchez-Martín et al., 2015; Canales et al.,2019). These cultivars have also been previously characterised by G x E interaction studies in field (Rispail et al., 2018; Sánchez-Martín et al., 2018; Sánchez-Martín et al., 2015). To our knowledge this is the first study that reveals the impact of drought on grain filling in oats. We have compared the metabolite profiles in several organs in these two oats cultivars to infer pathways that might explain differential strategies to cope with drought and grain filling performance.

### Drought induced source-sink dynamics suggests differential ‘drought escape’ mechanism in Patones and Flega

Mediterranean cultivars Flega and Patones exhibited differential responses to drought which could represent important adaptive mechanisms. The susceptible cultivar, Flega shows a ‘water saving’ strategy whereas the tolerant cultivar, Patones exhibits a ‘water spending’ strategy (Canales et al., 2021). Our observations confirm this observation, as compared to Patones, Flega showed reduced overall water usage (WUE) in both conditions **(Fig.1, Fig.S1).** In this study, these cultivars did not show symptoms of leaf curling, wilting, and yellowing with no significant changes in chlorophyll fluorescence during the drought treatment. There were some significant morphological parameters that differed between Patones and Flega with the latter showing faster flowering **(Fig.2)**. The morphological observation of grains showed an accelerated grain development in both cultivars, most prominently in Patones **(Fig.3)**. The observed morphological and grain size variation indicates that these two Mediterranean oat cultivars respond differently to drought. This led us to consider the metabolite responses in vegetative tissues and developing grains and how drought could impact on source-sink relationships.

Source-sink dynamics are very complex and is determined by multiple factors (Fabre et al., 2020), for example, biomass, relative CO_2_ levels and photosynthetic rates (Makino & Mae, 1999). Metabolically, the relative accumulation of photosynthetic end products (carbohydrates) in source tissues can reflect reduced sink demand and photosynthetic rates. This will also result in systemic decreases in nitrogen content (such as amino acids) in the leaves (Wei et al., 2019). Droughted Flega flag leaves appeared to show no compromised accumulation of glucose or glutamic acids suggesting that source strength from that organ was not altered. Further, the drought induced elevation in bioenergetic TCA intermediates such as citric acid **(Fig. 5)** which could be associated with the drought induced increased mobilisation of sucrose seen in the sheath **(Fig.5A)** and rachis **(Fig.5C)**. The lower levels of many amino acids (e. g. glutamine, lysine, asparagine, aspartic acid, but not tryptophan) in the sheath **(Fig.5A)** could reflect the demands of the sink tissue, as N influences the rate of grain filling and grain weight (Wei et al., 2019). In Patones, a drought induced reduction in glucose in the flag was one of the few significant changes in source tissues while amino acids changes were not prominent. Interestingly, an increase in raffinose family of oligosaccharides (RFOs) in the source tissues of both cultivars was observed **(Fig. 5 and 6)**. RFOs are a-1, 6- galactosyl extensions of sucrose that accumulate in seeds by phloem loading and transport and serve as desiccation protectants. This response seems to be accelerated by drought. Taken together our results suggests some over-lapping responses and also distinctive aspects of accelerated grain filling phenotype in the two cultivars.

### Metabolite profile shows accelerated grain filling in Patones that could be mediated by alpha-linolenic acid metabolism through jasmonate signaling

In our study showed that drought had no significant effect on spikelet numbers within the cultivars but were significant differences between cultivars. This suggests that any putative escape mechanism was primarily focused on grain development/ filling rate and not in the spikelet number. This highlighted the importance of a metabolomic approach to investigate the impact of drought on source-sink relationships.

Fatty acids (FAs) and lipids are essential components that also influence responses to abiotic and biotic stresses (He & Ding, 2020). In plants, the most predominant unsaturated fatty acids (UFAs) are 18-carbon (C18) including oleate, linoleate, and α-linoleate, which can act as antioxidants and also feed in the production of the important jasmonate (JAs) group of hormones (He & Ding, 2020). JAs include the oxygenated polyunsaturated fatty acids, ‘oxylipins’, JA and its 12-OPDA intermediate (Savchenko & Dehesh, 2014). Numerous studies have shown the role of JA in regulating abiotic and biotic stress responses. In this study we have observed changes in linolenic acid (a precursor for JA biosynthesis) and JA derivatives accumulation that may be related to drought stress and could be related to accelerated grain development in oat. Sánchez-Martín et al., (2018) noted that drought elicited linolenic acid processing leading to the accumulation of JAs in Patones which were not seen in Flega. Similarly, we observed a consistent increase in α-linolenic acid in the vegetative tissues of Patones which were not prominent in Flega **(Fig. 6D)**. Interestingly, 12-OPDA content in all the three source tissues of Patones, (PD-S, PD-FL, PD-R) did not show significant change but were lower in grains under drought. Reduced levels of 12-OPDA during drought could suggest increased flux towards jasmonic acid (JA) biosynthesis. Given the results we hypothesize a possible role of FAs and α-linolenic acid in particular, to increase JA signalling from sink to source (Dave & Graham, 2012). JAs and its derivatives are lipid-derived hormones known to contribute towards grain development and yield (Kim et al., 2009). Transgenic rice plants overexpressing the Arabidopsis JA carboxyl methyltransferase gene (*AtJMT*) with high levels of methyl jasmonate (MeJA) showed significant reduction in grain yield when exposed to drought stress (Kim et al., 2009). Recently, the wheat *triticale grain weight 1* (*tgw1*) mutant that has reduced grain weight has been shown to encode keto-acyl thiolase 2B (KAT-2B), which is involved in a peroxisomal step in JA biosynthesis (Chen et al., 2020). However, the *tgw1* mutation also compromised expression of the gibberellin biosynthesis gene, *ent*-kaurene synthase. In this context, it should be noted that we observed a significant accumulation of JA derivatives, and indeed, gibberellin in the grains of Patones, but not in Flega grains subjected to drought (**Fig.7,8**). Further work is crucial to dissect the interplay of hormones that influences grain development and yield during drought stress.

### Flega shows an oxidative stress under drought

A common observation in both cultivars was a shift in GSH/GSSG status, most likely reflecting oxidative stress because of drought. In Patones low levels of reduced glutathione was only seen in the sheath **(Fig. 6A)** whereas in Flega it was observed in all source tissues (**Fig. 5**) but not in grains (**Fig. 8**). The occurrence of high levels of ophthalmic acid (OA) also indicates that pronounced oxidative stress in many tissues is a feature in Flega. OA is synthesised from glutamate and 2-ABA in oat, barley, wheat, and rye (Servillo et al., 2018) and increase in OA is paralleled by decrease in GSH and increase in isoleucine biosynthesis. The generation of reactive oxygen species (ROS) can degrade lipids, proteins and carbohydrates resulting a shift in oxidative status of the cell (Kutasy et al., 2021). Grains of Flega in control and drought-treated plants exhibited similar metabolite profiles, although water deficit conditions resulted in increased accumulation of myo-inositol and TCA/glycolysis intermediates **(Fig.8).** This might suggest change in energy status which could be associated with oxidative stress and ROS accumulation. Oxidative stress can accelerate earlier flowering with oxidative stress and this has been extensively characterised in Arabidopsis. This model species was used to show how OXIDATIVE STRESS 2 effects on florigen FT and transcription factor FD (FLOWERING LOCUS D) (Blanvillain et al., 2011; Liang and Ow, 2019) and drought responsive changes in flowering acting via OXIDATIVE STRESS 3 flowering through APETALA 1 (Liang et al., 2019). Whilst, drought did not appear to accelerate flowering time in Flega, this mechanism (or similar) could be relevant here.

Equally, the signaling molecule myo-inositol phosphatase (MIPS) (Du et al., 2011) could be contributing to accelerated flowering in Flega. MIPS is associated with numerous stress responses and phosphoinositide (PI) signaling (Sharma et al., 2020) a role for this signal needs to be investigated in oats.

## Conclusion

This is the first report to elucidate the grain filling responses in oat under drought using non-targeted metabolite profiling. Our study on two Mediterranean oat cultivars Flega and Patones showed, to different extents, accelerated grain development in response to drought. Each cultivar showed significant differences in the drought associated shifts in the metabolomes of source and sink tissues. This suggest that stress associated changes in grain filling can occur through different/independent mechanisms by modulating source-sink activity. Patones showed differential accumulation of C18 FA’s and JA derivatives in source organs and grains under drought, and this may be linked to JA induced signaling responses. However, in Flega, we observed a shift in the central energy metabolism along with accumulation of oxidative markers in both source and sink organs and is likely to be linked to an accelerated flowering phenotype. These different mechanisms are shown schematically in Figure 9. Further dissection of these mechanisms with such as transcriptome studies could provide important information on central regulators that control accelerated grain filling in oat and that can contribute to better grain traits in drought-prone environments.

**Figure 9.**
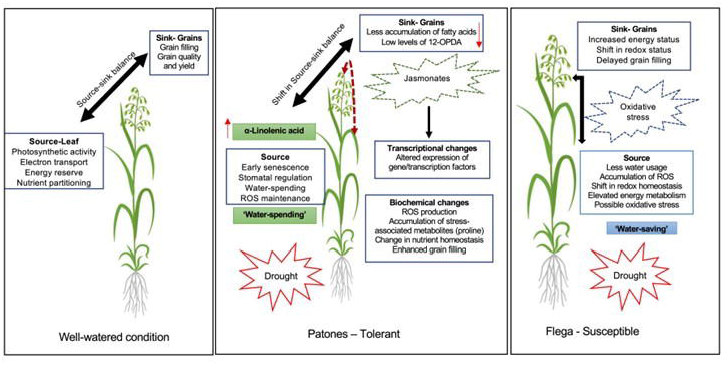
Proposed drought mechanism in Patones and Flega during grain filling. Schematic hypothetical representation of drought escape mechanism in Patones and Flega under drought during grain filling.

## Supplementary material

The following supplementary files are available at JXB online.

Fig. S1 (A) Patones and Flega at 75% SWC (control-left) and 25% SWC (drought-right). (B) The primary panicle was harvested at growth stage (GS)75. (C) The primary panicle inflorescence is divided into separate whorls. Spikelet differentiation occurs at the tip (“1”) and proceeds basally (sequentially “2”,”3” and “4”), and within spikelet floret develop acropetally. The top two spikelets from apical or top (1,2) and basal or bottom whorls (3,4) were harvested. The experiment was conducted on a Lematec platform at the National Plant Phenomics Centre (NPPC) under controlled conditions.

Fig. S2 Morphological parameters measured in Patones and Flega under control and drought treatments (A) Total plant weight (B) Stem height (C) Total panicle weight (D) Number of tillers. The data shows average ± SD, n=8 and statistically significance by T-test are denoted by small letters, *P ≤ 0.05*.

Fig. S3 PCA showing metabolite variation of all *m/z* features captured in negative mode (A) Top panel shows metabolite distribution (A) in Flega - source tissues and (B) in Flega - grains (C) Patones - source and (D) Patones - grains under well-watered and drought conditions.

Table S1 The annotated list of metabolites is provided in the table.

## Acknowledgements

We are grateful to Helen Philips for providing technical support to metabolite studies.

## Author contributions

LM, JD, and AG designed the study. AG, FC and JB conducted and monitored the experiments at NPPC; FC, JH, JB and KW imaging and data collection from Lemnatec phenomics platform. KW carried out phenotype analysis. AG, FJC, BH and RD conducted morphological, physiological measurements, grain imaging and metabolite sampling. AG and MB contributed to metabolite data analysis. FJC contributed to morphological and physiological and statistical aspects of the work. AG, FJC, EP, LM, JD and FC interpreted the results and drafted the manuscript. All authors participated in the critical reading and writing of the manuscript.

## Data Availability

Data supporting the findings in this study are available within the paper and supplementary data published online.

